# Spatiotemporal and demographic effects on avian malaria prevalence in blue tits

**DOI:** 10.64898/2026.04.15.718266

**Authors:** Angela Nicole Theodosopoulos, Fredrik Andreasson, Jane Jönsson, Johan Nilsson, Andreas Nord, Lars Råberg, Martin Stjernman, Ana Sofía Torres Lara, Jan-Åke Nilsson, Olof Hellgren

## Abstract

While the ubiquity of parasites is well understood, the extent to which host demographic factors shape parasite prevalence patterns merits further investigation. Using 15 years of breeding data from blue tits (*Cyanistes caeruleus*) in southern Sweden, spanning a 26-year timeframe, we assessed the roles of host age, sex, and field site on the odds of infection with three avian malaria parasite genera: *Haemoproteus, Plasmodium*, and *Leucocytozoon*. Further, we also evaluated the effects of these demographics on the odds of triple-genus coinfections. We found first-year breeders have fewer infections with *Haemoproteus* and *Plasmodium*, and fewer triple infections compared to older age classes, suggesting that birds are accumulating infections over time. *Leucocytozoon* infections are more prevalent in males than in females, and this may be due to sex-specific differences in physiology. The prevalence of malaria parasites and their coinfections also vary between the three sampling sites, indicative of an effect of host breeding habitat, even at a relatively small spatial scale (neighboring sites were separated by <5km). Across all three field sites, prevalence is overall significantly increasing over time. For *Haemoproteus*, this increase is more pronounced in older birds compared to younger birds. Such temporal changes in age-related infection patterns would not have been apparent without long-term data thus highlighting the importance of long-term studies for informing our understanding of host demographic effects on parasite prevalence.

## Introduction

Dependence for survival and energy acquisition of one organism (the parasite) at the expense of survival or reproductive success of another organism (the host) is one of the most common strategies in nature (Lafferty et al. 2008). Belonging to a highly ubiquitous and diverse group of organisms, parasites can dramatically shape their host populations, often while remaining hidden in plain sight (Anderson and May 1982).

Arguably, all metazoans are universally vulnerable to a set of parasites that they cannot escape (Poulin and Morand 2000), with parasite-imposed consequences ranging from relatively benign (Bensch et al. 2007), to severe effects on host survival (Van Riper et al. 1986, Coltman et al. 1999), and reproductive success (Marzal et al. 2005, Patterson et al. 2013). Parasite diversity in birds, and their likely influence on host fitness, has been explored across numerous systems. Among these studies, avian malaria parasites (Order: Haemosporida) are some of the most well-documented (Valkiūnas 2005); probably due to their cosmopolitan nature (Clark et al. 2014) and, in some cases, their relatively high prevalence (Greiner et al. 1975, Theodosopoulos et al. 2025). The genera *Haemoproteus, Plasmodium*, and *Leucocytozoon* are the most commonly studied avian malaria parasites (Valkiūnas 2005). Members of all three genera rely on separate vectors (e.g., biting midges: *Haemoproteus*, mosquitoes: *Plasmodium*, and *black flies*: *Leucocytozoon*) to transmit them to an avian host, where they asexually reproduce (Valkiūnas 2005).

Avian malaria infections can impose physiological consequences (Schoenle et al. 2017), accelerate aging through telomere shortening (Asghar et al. 2016), and reduce reproductive success (Knowles et al. 2010). The most well-known case of the negative impacts of avian malaria parasites is the introduction of *P. relictum* to the Hawaiian Islands, which is believed to have taken place in the early 1900’s (Van Riper et al. 1986). Most endemic bird species experienced severe population declines, and it is likely that some extinction events were the result of malarial disease (Warner 1968). However, malaria infections documented in wildlife are often chronic, where fitness effects are not directly observable (Bensch et al. 2007, Hammers et al. 2016).

Coinfections with more than one avian malaria parasite add an additional layer of complexity to these study systems. For instance, exposure to one parasite may facilitate subsequent infection by another, as was experimentally demonstrated for infections with *Plasmodium elongatum* that were facilitated by *Plasmodium relictum* (Palinauskas et al. 2018). Alternatively, interactions among parasite lineages may limit the establishment of secondary infections, resulting in fewer coinfections than expected. For example, a study of passerines in New Caledonia found negative associations between some malaria parasite lineages, suggestive of competitive interactions (Clark et al. 2016). Experiments are critical to explicitly test parasite facilitation vs. competition. However, if hosts accumulate infections over time, this pattern may reflect either facilitative interactions among parasites or largely neutral interactions in which multiple parasites can persist concurrently. Importantly, coinfections have been shown to impose additive costs on host fitness that exceed the consequences of individual infections (Marzal et al. 2008); thus, their dynamics merit further investigation.

An outstanding knowledge gap is what ecological and evolutionary mechanisms influence avian malaria prevalence. A good starting point is understanding how basic demographics such as host age, sex, and sampling location, correlate to parasite infections. The host breeding season is an ideal time to target these investigations given the heightened physiological stressors from reproduction, and the emergence of vectors during spring and summer (Beaudoin et al. 1971, Murdock et al. 2013). Elevated levels of testosterone in males may enhance the establishment of parasites (Folstad and Karter 1992, Deviche et al. 2001). However, multiple previous studies have found that females appear to have an overall higher prevalence of *Haemoproteus* compared to males (McCurdy et al. 1998). Spatially, birds tend to remain in small territories during the breeding season, and vectors, such as *Culicoides* biting midges are known to enter nest cavities enabling easier access to brooding females and nestlings (Žiegytė et al. 2021). As such, the habitat surrounding nesting sites can be an important predictor of these infections (Lutz et al. 2015). Additionally, age can shape malaria parasite abundance where older birds may have experienced more exposure events (Allander and Bennett 1994). However, nestlings and recently fledged birds may lack a well-developed immune system (Palacios et al. 2009), and this could lead to relatively younger birds having a higher infection odds following an exposure event.

We investigated the effects of host age, sex, and sampling site on avian malaria prevalence in a population of blue tits (*Cyanistes caeruleus*) by harnessing infection data from 15 breeding seasons spanning a 26-year study period for breeding blue tits near the Stensoffa Research Station in Southern Sweden (Figure 1, centered at approximately 55°42’17”N 13°26’48”E). Using data from three sampling sites, each separated by a distance of at least 5km, we screened birds for infections with *Haemoproteus, Plasmodium*, and *Leucocytozoon*, and their combined triple infections. In addition to these demographic factors, we also compared prevalence trends over time and across our sampling sites. We did so because a previous study found infections with all three malaria genera temporally increased at one of the field sites (Theodosopoulos et al. 2025), and we sought to explore the generality of this finding.

**Figure 1.**
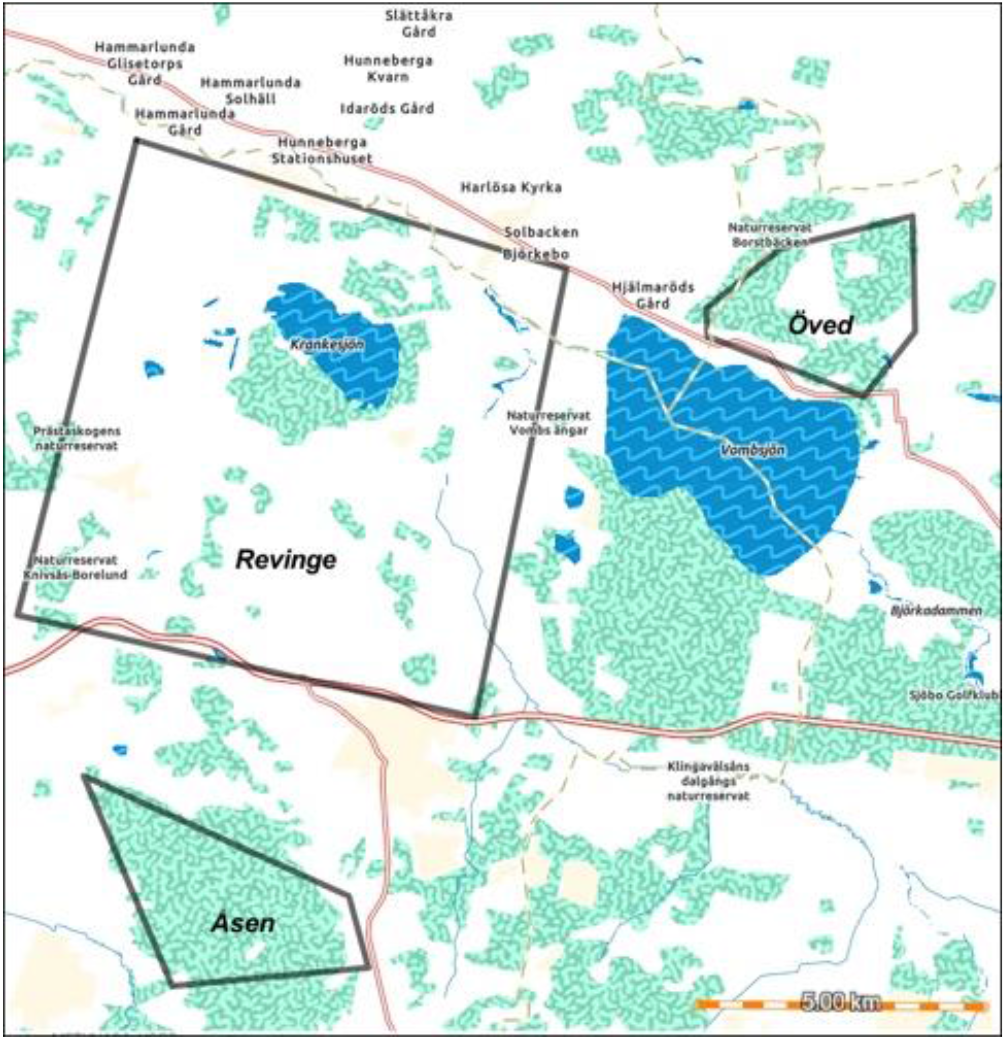
2498 samples were collected from blue tits at three sampling sites from 607 nest boxes between 1996 and 2021. Of these sampling events, 169 were from Åsen (sampled from 2007 – 2018), 364 were from Öved (sampled from 2007 – 2021), and 1965 were from Revinge (sampled from 1996 – 2021). For mapmaking, we used ArcGIS online with the ESRI enhanced contrast basemap.

## Material and methods

### Study system

Blue tits are small, partial-migrant passerines that nest in cavities, and studying their breeding is feasible with the use of artificial nesting snags or nest boxes (Svensson 1992). Prior to breeding, males establish and defend territories within which females build a cup-shaped nest using moss and other materials such as feathers (Stenning 2018). Females incubate a clutch ranging from 7 to 15 eggs (Nur 1986, Nilsson and Svensson 1993) and males often feed females during incubation (Nilsson and Smith 1988).

Between 1996 and 2021 we collected blood samples from breeding blue tits (*n* = 2498 sampling events) during the following years: 1996 – 2000, 2007 – 2011, and 2017 – 2021. Samples were collected from birds at three field sites (referred to as: Åsen, Öved, and Revinge, Figure 1), and from *n* = 607 different nest boxes. Of all sites in the study, Revinge had the most complete sampling (birds were sampled from Revinge during all 15 breeding seasons). Sampling at Öved and Åsen only took place after the year 2000 and we therefore lack data for earlier years. Additionally, we lack data for Åsen during later years of the study (2019 – 2021). Of the combined sampling events, *n* = 274 were from recaptured birds. None of these recaptures were sampled more than once during a single breeding season, and the maximum number of recapture events for any individual was four field seasons (Supplementary Figure 1). The breeding season took place between April and July, when females are easy to distinguish from males in the hand based on the presence of a brood patch (Svensson 1992). The overall sampling of blue tit males and females was highly similar between years and sampling sites (Supplementary Table 1). All birds were aged as (1) first-year breeders (henceforth ‘younger’), and (2) all older age classes (henceforth ‘older’) based on the presence of a molt limit between the primary and secondary coverts (indicating a younger bird; Svensson, 1992).

### Parasite screening

For all samples we conducted DNA extractions using previously described methods (Theodosopoulos et al. 2025). We used multiplex PCR (Ciloglu et al. 2019) to screen all birds for the presence of each of the three malaria genera. Samples were screened in duplicate given that some low intensity infections are not always detectable from a single round of screening (Ciloglu et al. 2019), and final scores were based on both screens. Additionally, we used negative controls, and positive controls for each of the three parasite genera. Positive infections were determined using gel electrophoresis where visualizing bands for the expected DNA fragment length determined the presence or absence of an infection with each parasite genus.

We used Sanger sequencing of *cytb* to type genetic lineages for a subset of *n* = 772 samples (Hellgren et al. 2004). *Haemoproteus* and *Plasmodium* were coamplified using the same two sets of primers and a separate set of primers was used to amplify *Leucocytozoon* (Hellgren et al. 2004). For *Haemoproteus* and *Plasmodium*, we typed samples from all 15 breeding seasons, and we typed samples from 1996 and 2020 – 2021 for *Leucocytozoon*. We screened samples twice and included both negative and positive controls. Sequences were edited using Geneious Prime software (v2025.0.3; Kearse et al. 2012) and aligned to the MalAvi database (Bensch et al. 2009) using the BLAST algorithm (Johnson et al. 2008) to determine lineage.

### Statistical analyses

Given the exploratory nature of our study, we used a multi-model comparison approach to assess the effects of demographic variables on the prevalence of infections with each parasite genus and of triple infections involving all three genera over time. All statistical analyses were conducted using R statistical software version 2024.12.1+563 (R Core Team 2013). We used the R package lme4 (Bates et al. 2003) to fit generalized linear mixed models (GLMMs) that assess how host age, sex, mean-centered sampling year, and their interactions influenced parasite prevalence. We also included host recapture status as a fixed effect to evaluate whether capture status is related to infection status. All models were binomial with a logit link function, and we included nest box as a random effect given that birds sampled from the same box may experience shared environmental conditions. We ran initial models based only on Revinge observations and fit a second set of models where sampling site was added as a fixed effect. Additionally, we set Revinge as the reference sampling site, meaning model estimates for Åsen and Öved were in comparison to Revinge. Models were fit using the bobyqa optimizer to aid convergence.

We used the R package MuMIn (Bartoń 2010) function *dredge* for model selection. Because *dredge* requires complete data for all predictors, we removed observations with missing values prior to analysis, resulting in 36 observations being dropped from the full dataset. The *dredge* output generates all possible sub-models and reports their Akaike Information Criterion corrected for small sample size (AICc) and associated model weights. Following previous studies, (Ferraguti et al. 2018, Rivero De Aguilar et al. 2018), we retained the top models defined by ≤ 2 AIC of the best model (Supplementary Table 2). We then applied model averaging using the *model.avg* function, and estimated predictor effects based on full-model averages (Table 1). We then evaluated model assumptions using the R package DHARMa (Hartig 2024) and simulated standardized residuals (1000 simulations per model), and used a combination of QQ plots and DHARMa diagnostic tests to examine residual uniformity, dispersion, zero inflation, and the presence of outliers (Supplementary Figures 2 & 3).

**Table 1.** Predictors for *Haemoproteus, Plasmodium*, and *Leucocytozoon* from top performing models (ΔAIC ≤ 2) fit to the “all sites” dataset. Model-averaged logistic regression coefficients on the log-odds scale (Log-odds), associated standard errors (SE), and p-values (p) are shown. Summed Akaike weight across the top-performing models in which each predictor was included.

| Parasite | Predictor | Log-odds | SE | p | Weight |
| --- | --- | --- | --- | --- | --- |
| *Haemoproteus* | (Intercept) | 1.36 | 0.09 | **< 2.2E-16** | NA |
| *Haemoproteus* | Age = Older | 1.14 | 0.15 | **< 2.2E-16** | 1 |
| *Haemoproteus* | Recaptured = TRUE | 0.26 | 0.28 | 3.43E-01 | 0.66 |
| *Haemoproteus* | Site = Åsen | -0.47 | 0.23 | **4.29E-02** | NA |
| *Haemoproteus* | Site = Öved | -1.24 | 0.15 | **< 2.2E-16** | NA |
| *Haemoproteus* | Year | 0.10 | 0.01 | **< 2.2E-16** | 1 |
| *Haemoproteus* | Age = Older:Year | 0.04 | 0.02 | **1.26E-02** | 1 |
| *Haemoproteus* | Sex = Male | 0.02 | 0.06 | 7.84E-01 | 0.21 |
| *Plasmodium* | (Intercept) | 0.70 | 0.08 | **< 2.2E-16** | NA |
| *Plasmodium* | Age = Older | 0.84 | 0.11 | **< 2.2E-16** | 1 |
| *Plasmodium* | Sex = Male | 0.09 | 0.10 | 3.85E-01 | 0.63 |
| *Plasmodium* | Site = Åsen | -0.42 | 0.20 | **3.54E-02** | NA |
| *Plasmodium* | Site = Öved | -1.16 | 0.14 | **< 2.2E-16** | NA |
| *Plasmodium* | Year | 0.06 | 0.01 | **< 2.2E-16** | 1 |
| *Plasmodium* | Recaptured = TRUE | -0.02 | 0.09 | 8.15E-01 | 0.18 |
| *Plasmodium* | Age = Older:Year | 0.00 | 0.01 | 8.25E-01 | 0.18 |
| *Leucocytozoon* | (Intercept) | 1.32 | 0.10 | **< 2.2E-16** | NA |
| *Leucocytozoon* | Age = Older | 0.14 | 0.15 | 3.23E-01 | 0.75 |
| *Leucocytozoon* | Sex = Male | 0.24 | 0.12 | 4.14E-02 | 1 |
| *Leucocytozoon* | Year | 0.03 | 0.01 | **6.17E-05** | 1 |
| *Leucocytozoon* | Age = Older:Year | -0.01 | 0.01 | 5.92E-01 | 0.35 |
| *Leucocytozoon* | Recaptured = TRUE | 0.05 | 0.13 | 7.15E-01 | 0.28 |
| Triple infection | (Intercept) | -0.39 | 0.08 | **2.00E-06** | NA |
| Triple infection | Age = Older | 0.83 | 0.10 | **< 2.2E-16** | 1 |
| Triple infection | Sex = Male | 0.11 | 0.10 | 2.85E-01 | 0.73 |
| Triple infection | Site = Åsen | -0.46 | 0.18 | **9.91E-03** | NA |
| Triple infection | Site = Öved | -0.99 | 0.14 | **< 2.2E-16** | NA |
| Triple infection | Year | 0.08 | 0.01 | **< 2.2E-16** | 1 |
| Triple infection | Recaptured = TRUE | 0.03 | 0.10 | 7.26E-01 | 0.24 |
| Triple infection | Age = Older:Year | -0.00 | 0.01 | 8.45E-01 | 0.12 |

### Visualizing model predictions

We used the R package ggeffects (Lüdecke 2018) to plot marginal effects using the *ggpredict* function, with predictions illustrated on the original (uncentered) year scale. Marginal effects and their 95% confidence intervals were plotted for these demographic variables, in addition to mean prevalence values using the R package ggplot2 (Wickham 2011). For visualizing site-level prevalence patterns, we removed model predictions for Åsen and Öved during the years in which we lacked sampling data (Supplementary Table 1).

## Results

### Parasite lineages

*H. majoris* (lineage: PARUS1) was the only *Haemoproteus* lineage occurring in the dataset of sequenced samples, and the most abundant lineage of our study (*n* = 630). Our ability to type *Plasmodium* was limited given that many positive-testing birds had coinfections with both *Plasmodium* and *Haemoproteus* (the latter is typically higher in intensity; Waldenström et al. 2004), resulting in sequencing bias. However, we did identify three different *Plasmodium* lineages: the most common being BT7 and TURDUS1 (both belonging to the species *P. circumflexum*, Valkiūnas et al. 2022), and two birds were infected with SGS1 (*P. relictum*, Hellgren et al. 2013). From a subset of 110 samples, we typed eight different *Leucocytozoon* lineages (See Supplementary Table 3) and 32 infections consisting of at least two *Leucocytozoon* lineages.

### Effect of age on prevalence

Across all sites, older birds had higher odds of being infected with *Haemoproteus* than younger birds (Table 1, Figure 2A). This age difference increased modestly over time, as indicated by a significant Age x Year interaction, where the odds of infection rose slightly faster in older birds than in younger birds. Older birds also had higher odds of being infected with *Plasmodium* (Figure 2B) and harbor triple infections (Figure 2D). However, we did not find any significant effect of age on *Leucocytozoon* prevalence (Figure 2C). Results were broadly consistent between the Revinge-only dataset and the full dataset (Supplementary Table 4, Supplementary Figure 4A–D). While host recapture status was retained in some top-ranked models, its estimated effects were consistently weak.

**Figure 2.**
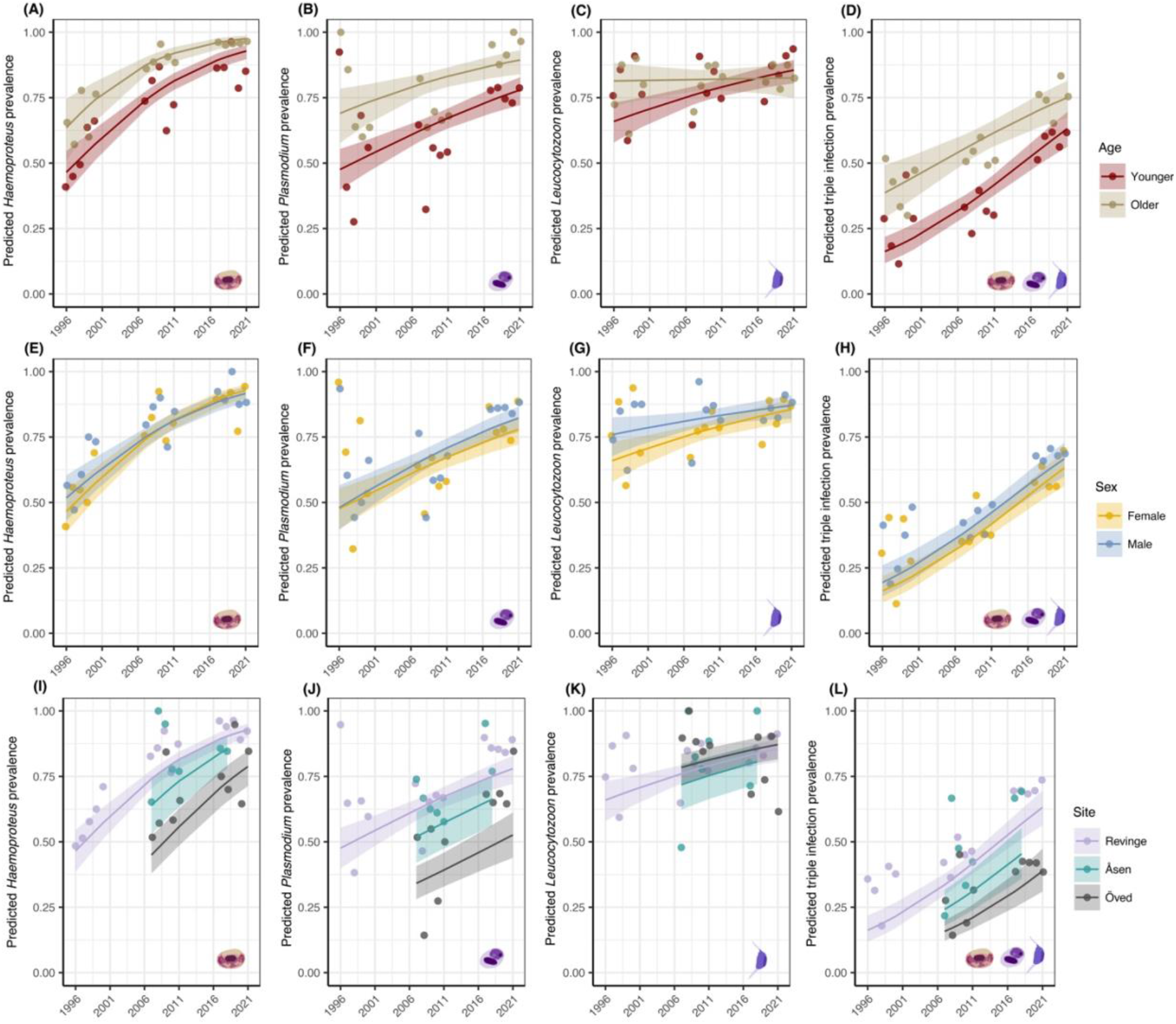
Estimated marginal effects (± 95% confidence intervals) for host demographic predictors of infection with *Haemoproteus, Plasmodium, Leucocytozoon*, and the occurrence of their combined triple infections. Plots show model-predicted effects of host age (A–D), sex (E–F), and sampling location (I–L). Age was categorized as first-year breeders (“younger”) versus all older age classes (“older”). Marginal effects and confidence intervals were generated using the *ggpredict* functions from the ggeffects R package (Lüdecke 2018). Points represent annual prevalence values and are overlaid for visualization using ggplot2 (Wickham 2011). Predictions are based on 2462 observations with complete data.

### Effect of sex on prevalence

Across all sites, males had higher odds of being infected with *Leucocytozoon* (Table 1, Figure 2G), and this pattern was consistent with results from the Revinge-only dataset (Supplementary Table 4, Supplementary Figure 4G). In contrast, sex was not a significant predictor for *Plasmodium* prevalence across all sites (Figure 2F), although males at Revinge were more likely to be infected than females (Supplementary Figure 4F). A similar pattern was observed for triple infections, with no significant sex effect across all sampling sites (Figure 2H), but males at Revinge had higher odds of harboring triple infections (Supplementary Figure 4H). There was no evidence of a sex-specific effect on *Haemoproteus* infection across all sites or within the Revinge-only dataset (Figure 2E; Supplementary Figure 4E).

### Effect of spatiotemporal variation on prevalence

Across all sites, the prevalence of all three malaria parasite genera increased significantly over time (Table 1, Figure 2I–L), consistent with findings from a previous study focused on temporal dynamics at Revinge (Theodosopoulos et al. 2025). The odds of infection increased steadily through time for each parasite genus, as well as for triple infections (Table 1). Temporal trends were similar between the Revinge-only and full datasets (Supplementary Table 4).

## Discussion

Across the study period, older birds consistently exhibited higher prevalence of *Haemoproteus* and *Plasmodium* infections, and a higher prevalence of triple infections compared to younger birds. Moreover, this age-related difference was not constant through time. For *Haemoproteus*, the odds of infection increased in both age classes, but rose slightly faster in older birds, resulting in a modest strengthening of age-related differences over time. Immunocompetence can be influenced by host age; for example, older mute swans (*Cygnus olor*) are more likely to immunologically responding to different viral subtypes of avian influenza relative to younger birds (Hill et al. 2016). However, in wild birds, avian malaria typically persists as chronic infections with low parasitemia (Valkiunas et al. 2004), suggesting that many infections are not fully cleared. The observed increased in prevalence with age suggests blue tits are accumulating *Haemoproteus* and *Plasmodium* infections over time. Similar findings have been reported in many other systems such as European bee-eaters (*Merops apiaster*; Emmenegger et al. 2020), purple-crowned fairy wrens (*Malurus coronatus*; Eastwood et al. 2019), and in other blue tit populations (Knowles et al. 2011, Podmokła et al. 2014). Importantly, increasing prevalence with age does not necessarily imply increasing parasite burden, as previous work in blue tits has shown that infection intensity may decline with host age, suggesting improved resistance through acquired immunity (Stjernman et al. 2004).

We did not observe a significant difference in *Leucocytozoon* prevalence between older and younger birds. One possible explanation is that hosts can clear *Leucocytozoon* infections, preventing an age-related accumulation of infections. If so, similar prevalences across age classes would be expected, consistent with our findings. However, blood-based detection might not reliably reflect long-term infection status given temporal variation in detectability and the inherent limitations of peripheral blood sampling. Because sampling was restricted to the breeding season, and the predicted climate window for *Leucocytozoon* transmission is from March 12^th^ – May 1^st^ (Theodosopoulos et al. 2025), our dataset may reflect a narrow temporal window in which *Leucocytozoon* infections are most likely detectable in blood.

Males were more likely to harbor *Leucocytozoon* infections than females, which may be partly explained by sex-specific hormonal differences (Folstad and Karter 1992). Particularly, the window of infection for this parasite coincides with the period of elevated testosterone levels in males due to establishing a breeding territory and attracting mates. We did not observe any sex-biased effects on *Haemoproteus* infections. In contrast to *Leucocytozoon, Haemoproteus* transmission likely occurs when birds are nestlings (Theodosopoulos et al. 2025). During the developmental stage, male and female nestlings do not differ in testosterone levels (Silverin and Sharp 1996), which may explain the lack of sex effects on *Haemoproteus* prevalence. Based on the Revinge-only dataset, males were more likely to be infected with *Plasmodium* than females, but this was not consistent with findings from the full dataset. This discrepancy may reflect unmeasured site-level covariates, or alternatively, the more intensive sampling at Revinge, which might have provided greater statistical power.

Blue tit populations at Åsen and Öved had significantly lower prevalence of *Haemoproteus* and *Plasmodium* compared to Revinge, and this could be due to habitat differences. Environmental differences in vegetation (Hernández-Lara et al. 2017), elevation (Theodosopoulos et al. 2023), and water quality (Okanga et al. 2013) are all described predictors of malaria parasite infections. The biting midges and mosquitoes that transmit *Haemoproteus* and *Plasmodium*, respectively, typically breed in standing water and moist grasslands (Uslu and Dik 2006, Fillinger et al. 2009, Burke et al. 2010, Noori et al. 2015). Blue tits at the Revinge field site had on average the most *Haemoproteus* and *Plasmodium* infections. Revinge is also the lowest elevation field site (∼25m) and surrounds a shallow lake along with encompassing ponds and bogs (Nilsson & Stjernman 2025). In contrast, Öved is the highest elevation sampling site (∼50m), and relatively dry compared to Åsen and Revinge. Previous work in the study system showed that biting midges were most abundant at Revinge (Nilsson & Stjernman 2025) suggesting that this site might provide more suitable breeding habitat for *Haemoproteus*-transmitting vectors. Nesting habitat may play a role in infection risk both at a broader scale (between field sites) and at a finer scale (within field sites).

Whether blue tits intentionally avoid nesting near suitable vector habitats remains unclear. However, previous studies have shown that some birds actively avoid nesting in places where there are ectoparasites (Bush and Clayton 2018), or brood parasites (Payne 1977, Forsman and Martin 2009). A more in-depth spatial analysis of blue tit nesting preferences correlated with their infection statuses could provide fruitful insight on this subject. We did not detect an effect of field site on *Leucocytozoon* prevalence. The black flies that transmit *Leucocytozoon*, commonly breed in flowing water (Adler and McCreadie 2019). Small streams are present across all three sampling locations, suggesting that suitable breeding habitat for black flies is broadly available across sites. Similarities in *Leucocytozoon* prevalence across sampling sites are unlikely to reflect frequent vector dispersal, as black flies typically disperse only short distances following emergence (<5 km; Adler and McCreadie 2019). Rather, the exposure risk is likely shaped by site-level vector population.

Across all sampling sites the prevalence of individual malaria parasite infections and triple infections increased over the 26-year study period (Figure 2, Figure 3). At Öved and Åsen, we observed patterns similar to what we previously reported for Revinge (Theodosopoulos et al. 2025). Importantly, while younger birds have fewer infections than older birds, and males have more *Leucocytozoon* infections than females, these patterns will probably become less apparent as prevalence increases in the overall population. The overall prevalence increase is likely not due to human-mediated landscape changes because all three field sites have remained relatively unaltered over the past 26 years. In a previous study, climate warming was found to drive the rise in malaria parasite prevalence at Revinge (Theodosopoulos et al. 2025), and the similar increase at nearby field sites might be indicative of a broader, regional pattern.

**Figure 3.**
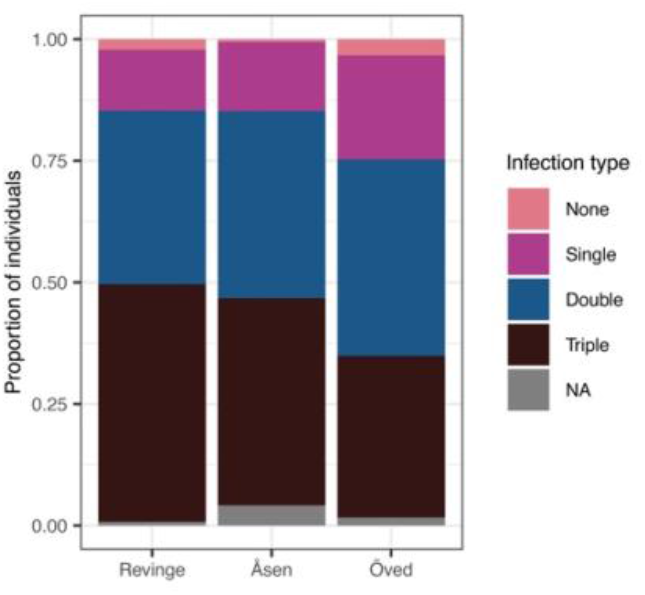
Stacked bar plot showing the proportion of individuals across infection categories for n = 2498 observations collected over the full study period. Bars represent the relative frequency of uninfected birds, birds with missing infection data (“NA”), and birds with single, double, and triple infections (i.e., infections with all three parasite genera), partitioned by sampling site.

## References

Adler, P. H. and McCreadie, J. W. 2019. Black Flies (Simuliidae). – Medical and Veterinary Entomology. Elsevier, pp. 237–259.

Allander, K. and Bennett, G. F. 1994. Prevalence and intensity of haematozoan infection in a population of great tits Parus major from Gotland, Sweden. – Journal of Avian Biology 25: 69.

Anderson, R. M. and May, R. M. 1982. Coevolution of hosts and parasites. – Parasitology 85: 411–426.

Asghar, M., Palinauskas, V., Zaghdoudi-Allan, N., Valkiūnas, G., Mukhin, A., Platonova, E., Färnert, A., Bensch, S. and Hasselquist, D. 2016. Parallel telomere shortening in multiple body tissues owing to malaria infection. – Proc. R. Soc. B. 283: 20161184.

Bartoń, K. 2025. Package ‘MuMIn’ v.1.48.11.

Bates, D., Maechler, M., Bolker, B. and Walker, S. 2003. Package ‘lme4’ v.1.1-37.

Beaudoin, R. L., Applegate, J. E., Davis, D. E. and McLEAN, R. G. 1971. A model for the ecology of avian malaria. – Journal of Wildlife Diseases 7: 5–13.

Bensch, S., Waldenström, J., Jonzén, N., Westerdahl, H., Hansson, B., Sejberg, D. and Hasselquist, D. 2007. Temporal dynamics and diversity of avian malaria parasites in a single host species. – Journal of Animal Ecology 76: 112–122.

Bensch, S., Hellgren, O. and Pérez-Tris, J. 2009. MalAvi: a public database of malaria parasites and related haemosporidians in avian hosts based on mitochondrial cytochrome b lineages. – Molecular Ecology Resources 9: 1353–1358.

Burke, R., Barrera, R., Lewis, M., Kluchinsky, T. and Claborn, D. 2010. Septic tanks as larval habitats for the mosquitoes Aedes aegypti and Culex quinquefasciatus in Playa-Playita, Puerto Rico. – Medical and Veterinary Entomology 24: 117–123.

Bush, S. E. and Clayton, D. H. 2018. Anti-parasite behaviour of birds. – Phil. Trans. R. Soc. B 373: 20170196.

Ciloglu, A., Ellis, V. A., Bernotienė, R., Valkiūnas, G. and Bensch, S. 2019. A new one-step multiplex PCR assay for simultaneous detection and identification of avian haemosporidian parasites. – Parasitol Res 118: 191–201.

Clark, N. J., Clegg, S. M. and Lima, M. R. 2014. A review of global diversity in avian haemosporidians (Plasmodium and Haemoproteus: Haemosporida): new insights from molecular data. – International Journal for Parasitology 44: 329–338.

Clark, N. J., Wells, K., Dimitrov, D. and Clegg, S. M. 2016. Co-infections and environmental conditions drive the distributions of blood parasites in wild birds. – Journal of Animal Ecology 85: 1461–1470.

Coltman, D. W., Pilkington, J. G., Smith, J. A. and Pemberton, J. M. 1999. Parasite-mediated selection against inbred Soay sheep in a free-living island population. – Evolution 53: 1259–1267.

Deviche, P., Greiner, E. C. and Manteca, X. 2001. Seasonal and age-related changes in blood parasite prevalence in dark-eye Juncos (Junco hyemalis, Aves, Passeriformes). – Journal of Experimental Zoology 289: 456–466.

Eastwood, J. R., Peacock, L., Hall, M. L., Roast, M., Murphy, S. A., Gonçalves Da Silva, A. and Peters, A. 2019. Persistent low avian malaria in a tropical species despite high community prevalence. – International Journal for Parasitology: Parasites and Wildlife 8: 88–93.

Emmenegger, T., Alves, J. A., Rocha, A. D., Costa, J. S., Schmid, R., Schulze, M. and Hahn, S. 2020. Population- and age-specific patterns of haemosporidian assemblages and infection levels in European bee-eaters (Merops apiaster). – International Journal for Parasitology 50: 1125–1131.

Ferraguti, M., Martínez-de La Puente, J., Bensch, S., Roiz, D., Ruiz, S., Viana, D. S., Soriguer, R. C. and Figuerola, J. 2018. Ecological determinants of avian malaria infections: An integrative analysis at landscape, mosquito and vertebrate community levels. – Journal of Animal Ecology 87: 727–740.

Fillinger, U., Sombroek, H., Majambere, S., Van Loon, E., Takken, W. and Lindsay, S. W. 2009. Identifying the most productive breeding sites for malaria mosquitoes in the Gambia. – Malar J 8: 62.

Folstad, I. and Karter, J. 1992. Parasites, bright males, and the immunocompetence handicap. – The American Naturalist 139: 603–622.

Forsman, J. T. and Martin, T. E. 2009. Habitat selection for parasite-free space by hosts of parasitic cowbirds. – Oikos 118: 464–470.

Greiner, E. C., Bennett, G. F., White, E. M. and Coombs, R. F. 1975. Distribution of the avian hematozoa of North America. – Can. J. Zool. 53: 1762–1787.

Hammers, M., Komdeur, J., Kingma, S. A., Hutchings, K., Fairfield, E. A., Gilroy, D. L. and Richardson, D. S. 2016. Age-specific haemosporidian infection dynamics and survival in Seychelles warblers. – Sci Rep 6: 29720.

Hartig, F. 2024. DHARMa: Residual diagnostics for hierarchical (multi-level/mixed) regression models. v0.4.7.

Hellgren, O., Waldenström, J. and Bensch, S. 2004. A new PCR assay for simultaneous studies of Leucocytozoon, Plasmodium, and Haemoproteus from avian blood. – Journal of Parasitology 90: 797–802.

Hellgren, O., Kutzer, M., Bensch, S., Valkiūnas, G. and Palinauskas, V. 2013. Identification and characterization of the merozoite surface protein 1 (msp1) gene in a host-generalist avian malaria parasite, Plasmodium relictum (lineages SGS1 and GRW4) with the use of blood transcriptome. – Malar J 12: 381.

Hernández-Lara, C., González-García, F. and Santiago-Alarcon, D. 2017. Spatial and seasonal variation of avian malaria infections in five different land use types within a neotropical montane forest matrix. – Landscape and Urban Planning 157: 151–160.

Hill, S. C., Manvell, R. J., Schulenburg, B., Shell, W., Wikramaratna, P. S., Perrins, C., Sheldon, B. C., Brown, I. H. and Pybus, O. G. 2016. Antibody responses to avian influenza viruses in wild birds broaden with age. – Proc. R. Soc. B. 283: 20162159.

Johnson, M., Zaretskaya, I., Raytselis, Y., Merezhuk, Y., McGinnis, S. and Madden, T. L. 2008. NCBI BLAST: a better web interface. – Nucleic Acids Research 36: W5–W9.

Kearse, M., Moir, R., Wilson, A., Stones-Havas, S., Cheung, M., Sturrock, S., Buxton, S., Cooper, A., Markowitz, S., Duran, C., Thierer, T., Ashton, B., Meintjes, P. and Drummond, A. 2012. Geneious Basic: An integrated and extendable desktop software platform for the organization and analysis of sequence data. – Bioinformatics 28: 1647–1649.

Knowles, S. C. L., Palinauskas, V. and Sheldon, B. C. 2010. Chronic malaria infections increase family inequalities and reduce parental fitness: experimental evidence from a wild bird population. – Journal of Evolutionary Biology 23: 557–569.

Knowles, S. C. L., Wood, M., Alves, R., Wilkin, T. A., Bensch, S. and Sheldon, B. C. 2011. Molecular epidemiology of malaria prevalence and parasitaemia in a wild bird population. – Molecular Ecology 20: 1062–1076.

Lafferty, K. D., Allesina, S., Arim, M., Briggs, C. J., De Leo, G., Dobson, A. P., Dunne, J. A., Johnson, P. T. J., Kuris, A. M., Marcogliese, D. J., Martinez, N. D., Memmott, J., Marquet, P. A., McLaughlin, J. P., Mordecai, E. A., Pascual, M., Poulin, R. and Thieltges, D. W. 2008. Parasites in food webs: the ultimate missing links. – Ecology Letters 11: 533–546.

Lüdecke, D. 2018. ggeffects: Tidy Data Frames of Marginal Effects from Regression Models. v.2.2.1

Lutz, H. L., Hochachka, W. M., Engel, J. I., Bell, J. A., Tkach, V. V., Bates, J. M., Hackett, S. J. and Weckstein, J. D. 2015. Parasite Prevalence Corresponds to Host Life History in a Diverse Assemblage of Afrotropical Birds and Haemosporidian Parasites. – PLoS ONE 10: e0121254.

Marzal, A., Lope, F. D., Navarro, C. and Mller, A. P. 2005. Malarial parasites decrease reproductive success: an experimental study in a passerine bird. – Oecologia 142: 541–545.

Marzal, A., Bensch, S., Reviriego, M., Balbontin, J. and De Lope, F. 2008. Effects of malaria double infection in birds: one plus one is not two. – J of Evolutionary Biology 21: 979–987.

McCurdy, D. G., Shutler, D., Mullie, A. and Forbes, M. R. 1998. Sex-biased parasitism of avian hosts: relations to blood parasite taxon and mating system. – Oikos 82: 303.

Murdock, C. C., Foufopoulos, J. and Simon, C. P. 2013. A transmission model for the ecology of an avian blood parasite in a temperate ecosystem. – PLoS ONE 8: e76126.

Nilsson, J.-Å. and Smith, H. G. 1988. Incubation feeding as a male tactic for early hatching. – Animal Behaviour 36: 641–647.

Nilsson, J.-A. and Svensson, E. 1993. Energy constraints and ultimate decisions during egg-laying in the blue tit. – Ecology 74: 244–251.

Nilsson, J.-A. and Stjernman, M. 2025. Spatial and temporal variation in biting midge (Culicoides) abundance in active nest boxes. – bioRxiv 2025.07.21.665931.

Noori, N., Lockaby, B. G. and Kalin, L. 2015. Larval development of Culex quinquefasciatus in water with low to moderate. – Journal of Vector Ecology 40: 208–220.

Nur, N. 1986. Is clutch size variation in the blue tit (Parus caeruleus) adaptive? An experimental study. – The Journal of Animal Ecology 55: 983.

Okanga, S., Cumming, G. S. and Hockey, P. A. 2013. Avian malaria prevalence and mosquito abundance in the Western Cape, South Africa. – Malar J 12: 370.

Palacios, M. G., Cunnick, J. E., Vleck, D. and Vleck, C. M. 2009. Ontogeny of innate and adaptive immune defense components in free-living tree swallows, Tachycineta bicolor. – Developmental & Comparative Immunology 33: 456–463.

Palinauskas, V., Žiegytė, R., Šengaut, J. and Bernotienė, R. 2018. Different paths – the same virulence: experimental study on avian single and co-infections with Plasmodium relictum and Plasmodium elongatum. – International Journal for Parasitology 48: 1089–1096.

Patterson, J. E. H., Neuhaus, P., Kutz, S. J. and Ruckstuhl, K. E. 2013. Parasite removal improves reproductive success of female North American red squirrels (Tamiasciurus hudsonicus). – PLoS ONE 8: e55779.

Payne, R. B. 1977. The ecology of brood parasitism in birds. – Annu. Rev. Ecol. Syst. 8: 1–28.

Podmokła, E., Dubiec, A., Drobniak, S. M., Arct, A., Gustafsson, L. and Cichoń, M. 2014. Determinants of prevalence and intensity of infection with malaria parasites in the Blue Tit. – J Ornithol 155: 721–727.

Poulin, R. and Morand, S. 2000. The diversity of parasites. – The Quarterly Review of Biology 75: 277–293.

R Core Team. 2013. R: A language and environment for statistical computing.

Rivero De Aguilar, J., Castillo, F., Moreno, A., Peñafiel, N., Browne, L., Walter, S. T., Karubian, J. and Bonaccorso, E. 2018. Patterns of avian haemosporidian infections vary with time, but not habitat, in a fragmented Neotropical landscape. – PLoS ONE 13: e0206493.

Schoenle, L. A., Kernbach, M., Haussmann, M. F., Bonier, F. and Moore, I. T. 2017. An experimental test of the physiological consequences of avian malaria infection. – Journal of Animal Ecology 86: 1483–1496.

Schlicht L., Valcu, M., Kempenaers, B. 2015. Male extraterritorial behavior predicts extrapair paternity pattern in blue tits, Cynaistes caeruleus. – Behavioral Ecology 26: 1404–1413.

Silverin, B. and Sharp, P. 1996. The development of the hypothalamic-pituitary-gonadal axis in juvenile great tits. – General and Comparative Endocrinology 103: 150–166.

Sol, D., Jovani, R. and Torres, J. 2003. Parasite mediated mortality and host immune response explain age-related differences in blood parasitism in birds. – Oecologia 135: 542–547.

Stenning, M. 2018. The blue tit. – Bloomsbury Publishing Plc.

Stjernman, M., Råberg, L. and Nilsson, J. 2004. Survival costs of reproduction in the blue tit (Parus caeruleus): a role for blood parasites? – Proc. R. Soc. Lond. B 271: 2387–2394.

Svensson, L. 1992. Identification guide to Euopean passerines. – Cornell University Press.

Theodosopoulos, A. N., Grabenstein, K. C., Larrieu, M. E., Arnold, V. and Taylor, S. A. 2023. Similar parasite communities but dissimilar infection patterns in two closely related chickadee species. – Ornithology 140: ukad033.

Theodosopoulos, A. N., Andreasson, F., Jönsson, J., Nilsson, J., Nord, A., Råberg, L., Stjernman, M., Lara, A. S. T., Nilsson, J. and Hellgren, O. 2025b. Climate-Driven Increase in Transmission of a Wildlife Malaria Parasite Over the Last Quarter Century. – Global Change Biology 31: e70550.

Uslu, U. and Dik, B. 2006. Vertical distribution of Culicoides larvae and pupae. – Medical Vet Entomology 20: 350–352.

Valkiūnas, G. 2005. Avian malaria and other haemosporidia. – CRC Press.

Valkiūnas, G., Duc, M. and Iezhova, T. A. 2022. Increase of avian Plasmodium circumflexum prevalence, but not of other malaria parasites and related haemosporidians in northern Europe during the past 40 years. – Malar J 21: 105.

Van Riper, C., Van Riper, S. G., Goff, M. L. and Laird, M. 1986. The Epizootiology and Ecological Significance of Malaria in Hawaiian Land Birds. – Ecological Monographs 56: 327–344.

Waldenström, J., Bensch, S., Hasselquist, D. and Östman, Ö. 2004. A New Nested Polymerase Chain Reaction Method Very Efficient in Detecting Plasmodium and Haemoproteus Infections From Avian Blood. – Journal of Parasitology 90: 191–194.

Warner, R. E. 1968. The Role of Introduced Diseases in the Extinction of the Endemic Hawaiian Avifauna. – The Condor 70: 101–120.

Wickham, H. 2011. ggplot2. – WIREs Computational Statistics 3: 180–185.

Žiegytė, R., Platonova, E., Kinderis, E., Mukhin, A., Palinauskas, V. and Bernotienė, R. 2021. Culicoides biting midges involved in transmission of haemoproteids. – Parasites Vectors 14: 27.

Zylberberg, M., Derryberry, E. P., Breuner, C. W., Macdougal-Shackletonl, E. A., Cornelius, J. M. and Hahn, T. P. 2015. <i>Haemoproteus<i> infected birds have increased lifetime reproductive success. – Parasitology 14: 1033–1043.

