## Supplementary Figures and Tables for "Spatiotemporal and demographic effects on avian malaria prevalence in blue tits"

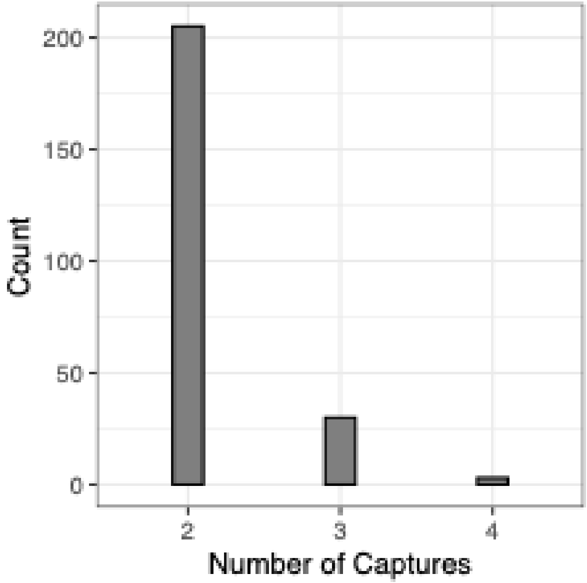


**Figure 1 –** Of the 2498 sampling events used for this study, 274 were form recaptured birds. At most, individual birds were captured four times, and most recaptured individuals were captured twice.

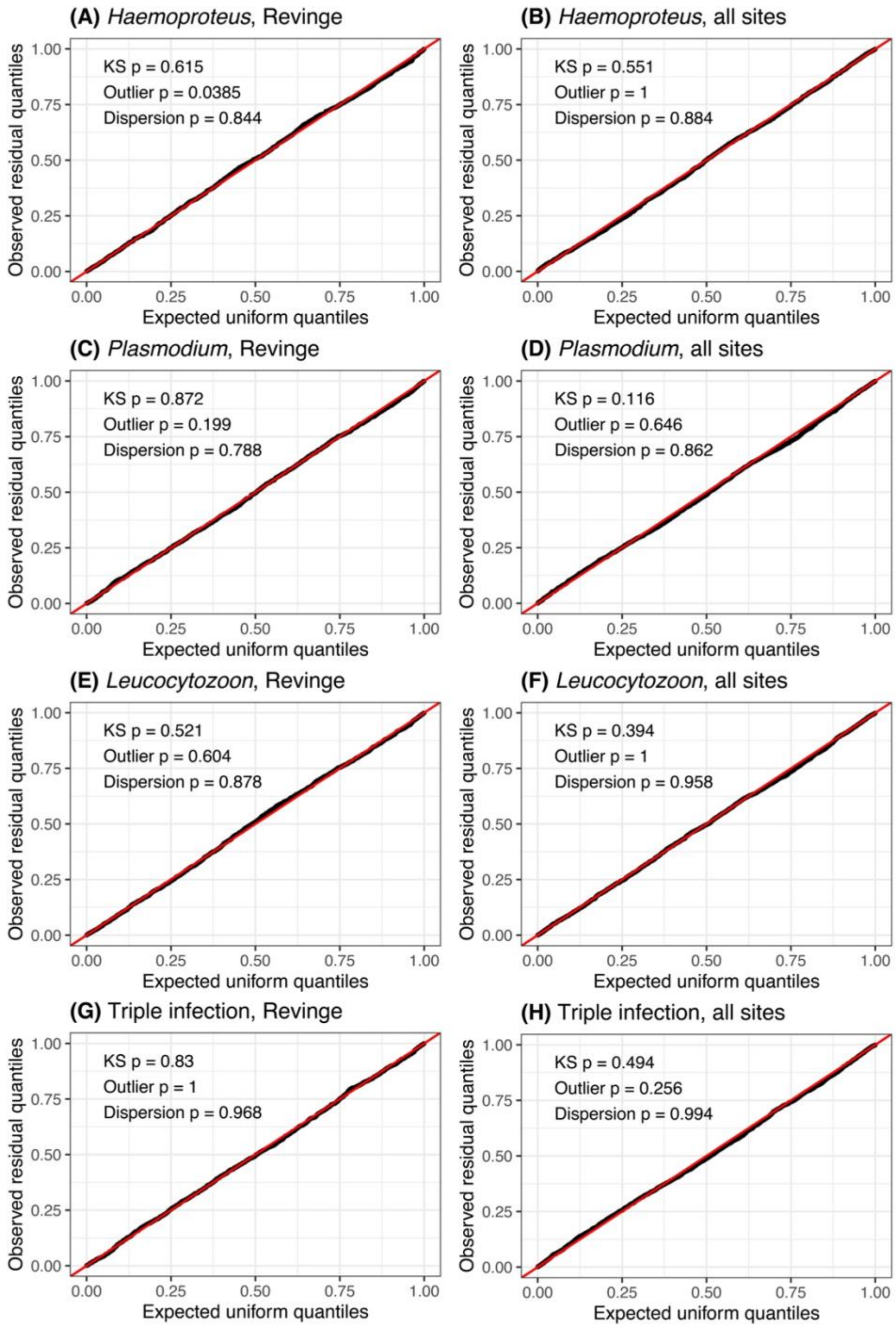


**Figure 2 –** Quantile-quantile (QQ) plots of scaled residuals generated using the R package DHARMa (Hartig 2024) for binomial mixed-effects models of *Haemoproteus* (A–B), *Plasmodium* (C–D), *Leucocytozoon* (E–F), and the presence of triple infections (G–H). For each parasite group, plots show results for Revinge-only data and data from all sites, respectively. Observed residual quantiles closely followed an expected uniform distribution and Kolmogorov-Smirnov (KS), outlier, and dispersion tests indicated generally good model fit with no substantial deviations from model assumptions.

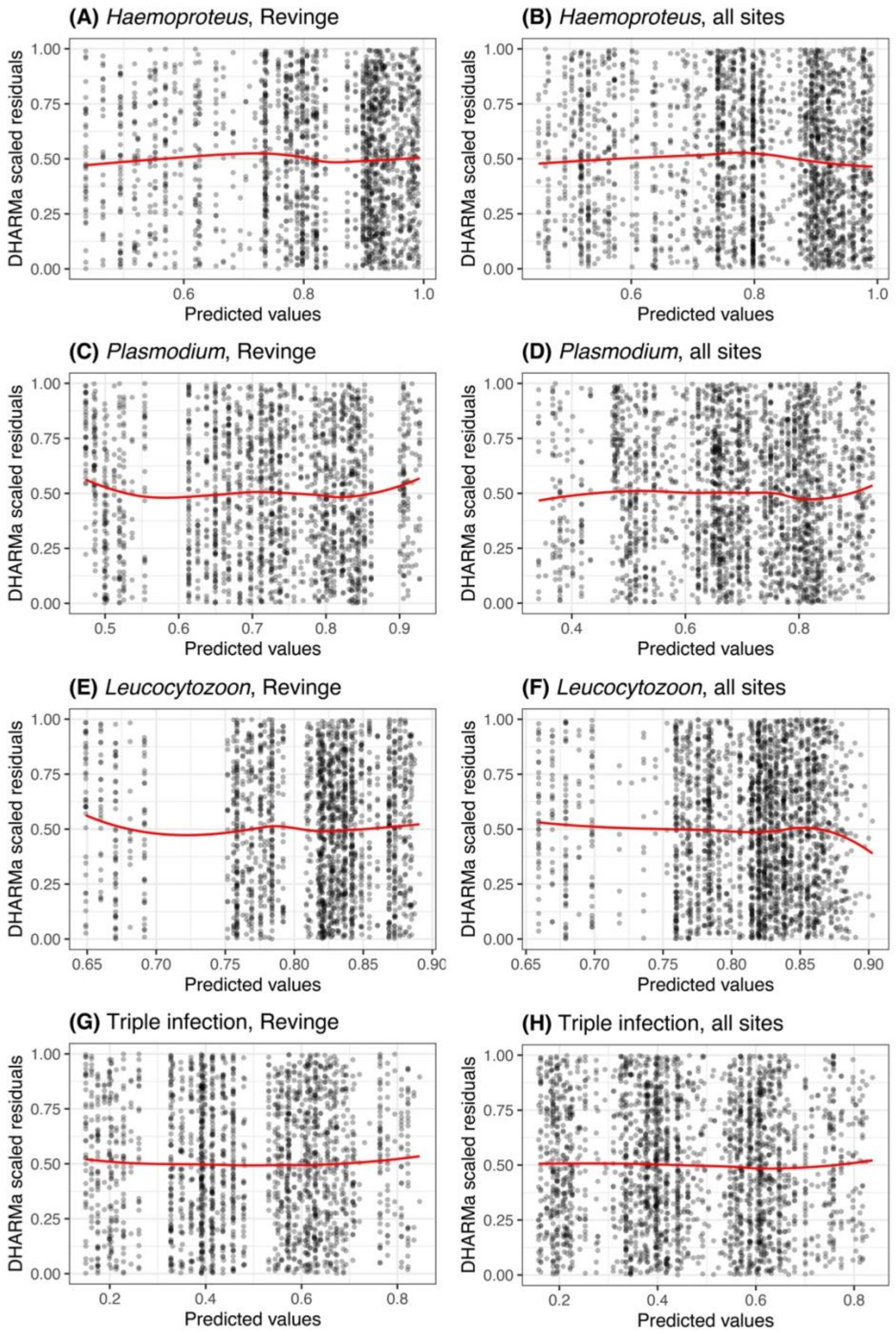


**Figure 3 –** Scaled DHARMa residuals from 1000 simulations based on fitted binomial mixed-effects models (refit = FALSE) using the R package DHARMa (Hartig 2024). Residuals are plotted against rank-transformed predicted values and red curves show LOESS-smoothing trends. Across models, residuals showed no strong systematic patterns across the range of predictors, indicating generally adequate model fits. Plots correspond to models for *Haemoproteus* (A–B), *Plasmodium* (C–D), *Leucocytozoon* (E–F), and the presence of triple infections (G–H), with analyses conducted using Revinge-only data and all sampling sites, respectively.

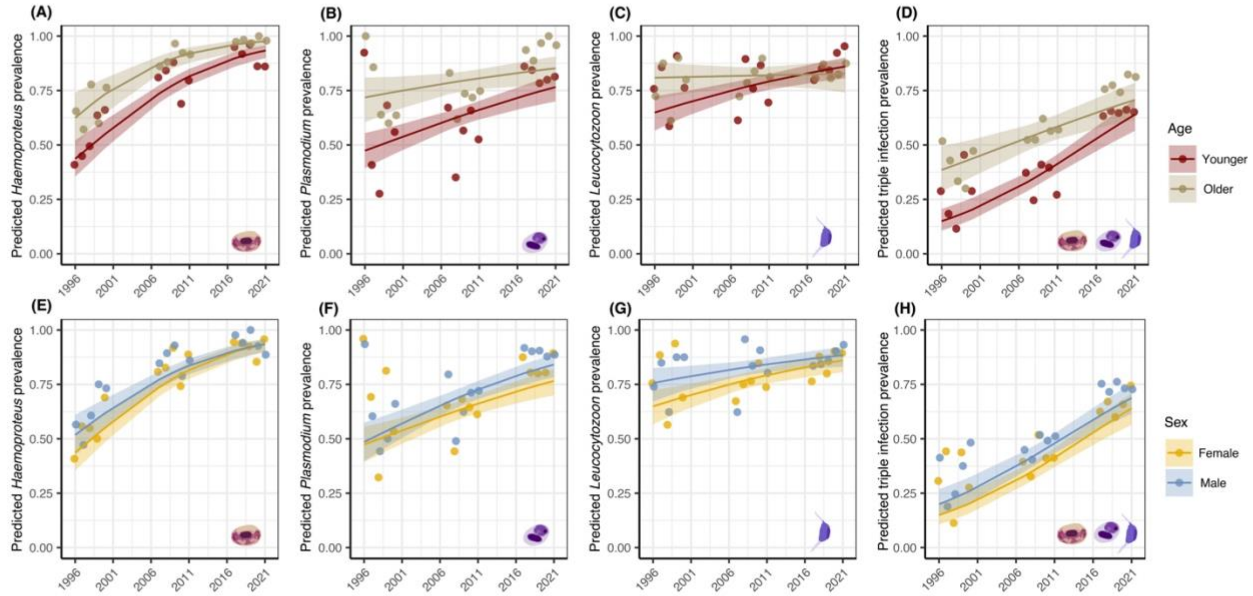


**Figure 4 –** Estimated marginal effects (± 95% confidence intervals) for host demographic predictors of infection with *Haemoproteus*, *Plasmodium*, *Leucocytozoon*, and the occurrence of their combined triple infections. Plots show model-predicted effects of host age (A–D) and sex (E–F). Age was categorized as first-year breeders ("younger") versus all older age classes ("older"). Marginal effects and confidence intervals were generated using the *ggpredict* functions from the ggeffects R package (Lüdecke 2018). Points represent annual prevalence values and are overlaid for visualization using ggplot2 (Wickham 2011). Predictions are based on a subset of n = 1944 at the Revinge field site with complete data.

**Table 1 —** Abundance of males, females, first-year breeders, and all older age classes spanning each sampling site and each year of the study.

| Site | Year | Site_year | Males | Females | First-year breeders | Older age classes |
|---|---|---|---|---|---|---|
| Asen | 1996 | Asen_1996 | 0 | 0 | 0 | 0 |
| Oved | 1996 | Oved_1996 | 0 | 0 | 0 | 0 |
| Revinge | 1996 | Revinge_1996 | 48 | 50 | 67 | 29 |
| Asen | 1997 | Asen_1997 | 0 | 0 | 0 | 0 |
| Oved | 1997 | Oved_1997 | 0 | 0 | 0 | 0 |
| Revinge | 1997 | Revinge_1997 | 53 | 52 | 49 | 56 |
| Asen | 1998 | Asen_1998 | 0 | 0 | 0 | 0 |
| Oved | 1998 | Oved_1998 | 0 | 0 | 0 | 0 |
| Revinge | 1998 | Revinge_1998 | 61 | 62 | 87 | 36 |
| Asen | 1999 | Asen_1999 | 0 | 0 | 0 | 0 |
| Oved | 1999 | Oved_1999 | 0 | 0 | 0 | 0 |
| Revinge | 1999 | Revinge_1999 | 16 | 16 | 22 | 10 |
| Asen | 2000 | Asen_2000 | 0 | 0 | 0 | 0 |
| Oved | 2000 | Oved_2000 | 0 | 0 | 0 | 0 |
| Revinge | 2000 | Revinge_2000 | 56 | 58 | 59 | 55 |
| Asen | 2007 | Asen_2007 | 11 | 12 | 14 | 9 |
| Oved | 2007 | Oved_2007 | 14 | 15 | 24 | 5 |
| Revinge | 2007 | Revinge_2007 | 98 | 105 | 137 | 65 |
| Asen | 2008 | Asen_2008 | 1 | 2 | 2 | 1 |
| Oved | 2008 | Oved_2008 | 4 | 3 | 6 | 1 |
| Revinge | 2008 | Revinge_2008 | 49 | 55 | 60 | 43 |
| Asen | 2009 | Asen_2009 | 20 | 20 | 17 | 23 |
| Oved | 2009 | Oved_2009 | 26 | 26 | 29 | 22 |
| Revinge | 2009 | Revinge_2009 | 85 | 85 | 83 | 87 |
| Asen | 2010 | Asen_2010 | 21 | 20 | 15 | 26 |
| Oved | 2010 | Oved_2010 | 44 | 46 | 63 | 26 |
| Revinge | 2010 | Revinge_2010 | 121 | 128 | 169 | 80 |
| Asen | 2011 | Asen_2011 | 14 | 14 | 7 | 21 |
| Oved | 2011 | Oved_2011 | 19 | 19 | 18 | 20 |
| Revinge | 2011 | Revinge_2011 | 87 | 80 | 59 | 108 |
| Asen | 2017 | Asen_2017 | 10 | 11 | 10 | 11 |
| Oved | 2017 | Oved_2017 | 23 | 21 | 28 | 16 |
| Revinge | 2017 | Revinge_2017 | 85 | 73 | 79 | 78 |
| Asen | 2018 | Asen_2018 | 7 | 6 | 9 | 4 |
| Oved | 2018 | Oved_2018 | 20 | 20 | 25 | 15 |
| Revinge | 2018 | Revinge_2018 | 102 | 82 | 122 | 62 |
| Asen | 2019 | Asen_2019 | 0 | 0 | 0 | 0 |
| Oved | 2019 | Oved_2019 | 9 | 11 | 5 | 15 |
| Revinge | 2019 | Revinge_2019 | 43 | 40 | 52 | 31 |
| Asen | 2020 | Asen_2020 | 0 | 0 | 0 | 0 |
| Oved | 2020 | Oved_2020 | 15 | 16 | 24 | 7 |
| Revinge | 2020 | Revinge_2020 | 41 | 41 | 65 | 17 |
| Asen | 2021 | Asen_2021 | 0 | 0 | 0 | 0 |
| Oved | 2021 | Oved_2021 | 7 | 6 | 4 | 9 |
| Revinge | 2021 | Revinge_2021 | 46 | 47 | 44 | 48 |
| | | **TOTAL** | 1256 | 1242 | 1454 | 1036 |

**Table 2 –** Summary of top-performing generalized linear mixed-effects models predicting the probability of infection with *Haemoproteus*, *Plasmodium*, *Leucocytozoon*, and the occurrence of triple infections. Models are shown for analyses restricted to the Revinge population and for analyses including all sampling sites. Model selection was based on Akaike's Information Criterion corrected for small sample size (AICc). For each response, all candidate models with ∆AIC ≤ 2 are reported, along with model degrees of freedom (Df), log-likelihood (LogLik), AICc, ∆AIC, and Akaike weight. Predictor columns indicate whether a term was included (+) or excluded (NA) from a given model, and intercept values are reported on the logit scale. Fixed effects considered included host age, sex, recapture status, mean-centered sampling year, and sampling site (for all-sites models). Interaction terms were evaluated among age, sex, and sampling year only.

| Model | Df | LogLik | AICc | Delta | Weight | (Intercept) | Age | Recap. | Sex | Year | Age:Sex | Age:Year | Sex:Year | Age:Sex:Year | Site |
|---|---|---|---|---|---|---|---|---|---|---|---|---|---|---|---|
| haem_revinge_model1 | 6 | -810.5 | 1633.0 | 0.00 | 0.16 | 1.35 | + | NA | + | 0.11 | NA | + | NA | NA | NA |
| haem_revinge_model2 | 5 | -811.7 | 1633.4 | 0.37 | 0.14 | 1.44 | + | NA | NA | 0.11 | NA | + | NA | NA | NA |
| haem_revinge_model3 | 7 | -809.9 | 1633.9 | 0.93 | 0.10 | 1.35 | + | + | + | 0.11 | NA | + | NA | NA | NA |
| haem_revinge_model4 | 6 | -811.1 | 1634.2 | 1.23 | 0.09 | 1.44 | + | + | NA | 0.11 | NA | + | NA | NA | NA |
| plas_revinge_model1 | 6 | -1092.9 | 2197.9 | 0.00 | 0.21 | 0.60 | + | NA | + | 0.05 | NA | NA | + | NA | NA |
| plas_revinge_model2 | 5 | -1094.4 | 2198.8 | 0.90 | 0.13 | 0.62 | + | NA | + | 0.06 | NA | NA | NA | NA | NA |
| plas_revinge_model3 | 7 | -1092.7 | 2199.5 | 1.66 | 0.09 | 0.60 | + | + | + | 0.05 | NA | NA | + | NA | NA |
| plas_revinge_model4 | 7 | -1092.8 | 2199.6 | 1.68 | 0.09 | 0.61 | + | NA | + | 0.05 | NA | + | + | NA | NA |
| plas_revinge_model5 | 7 | -1092.9 | 2199.9 | 2.00 | 0.08 | 0.61 | + | NA | + | 0.05 | + | NA | + | NA | NA |
| leuco_revinge_model1 | 4 | -958.8 | 1925.6 | 0.00 | 0.12 | 1.37 | NA | NA | + | 0.04 | NA | NA | NA | NA | NA |
| leuco_revinge_model2 | 5 | -957.9 | 1925.9 | 0.29 | 0.10 | 1.31 | + | NA | + | 0.04 | NA | NA | NA | NA | NA |
| leuco_revinge_model3 | 6 | -957.1 | 1926.3 | 0.67 | 0.08 | 1.33 | + | NA | + | 0.04 | NA | + | NA | NA | NA |
| leuco_revinge_model4 | 6 | -957.2 | 1926.4 | 0.80 | 0.08 | 1.26 | + | NA | + | 0.04 | + | NA | NA | NA | NA |
| leuco_revinge_model5 | 7 | -956.4 | 1926.8 | 1.20 | 0.07 | 1.27 | + | NA | + | 0.04 | + | + | NA | NA | NA |
| leuco_revinge_model6 | 5 | -958.5 | 1927.1 | 1.50 | 0.06 | 1.36 | NA | + | + | 0.04 | NA | NA | NA | NA | NA |
| leuco_revinge_model7 | 5 | -958.8 | 1927.6 | 2.00 | 0.04 | 1.37 | NA | NA | + | 0.03 | NA | NA | + | NA | NA |
| triple_revinge_model1 | 5 | -1227.5 | 2465.0 | 0.00 | 0.16 | -0.43 | + | NA | + | 0.08 | NA | NA | NA | NA | NA |
| triple_revinge_model2 | 6 | -1226.8 | 2465.6 | 0.56 | 0.12 | -0.43 | + | NA | + | 0.09 | NA | + | NA | NA | NA |
| triple_revinge_model3 | 6 | -1227.0 | 2466.1 | 1.06 | 0.09 | -0.43 | + | + | + | 0.08 | NA | NA | NA | NA | NA |
| triple_revinge_model4 | 6 | -1227.2 | 2466.3 | 1.34 | 0.08 | -0.44 | + | NA | + | 0.08 | NA | NA | + | NA | NA |
| triple_revinge_model5 | 7 | -1226.3 | 2466.6 | 1.56 | 0.07 | -0.43 | + | + | + | 0.09 | NA | + | NA | NA | NA |
| triple_revinge_model6 | 7 | -1226.3 | 2466.8 | 1.75 | 0.06 | -0.44 | + | NA | + | 0.08 | NA | + | + | NA | NA |
| triple_revinge_model7 | 6 | -1227.5 | 2467.0 | 1.97 | 0.06 | -0.44 | + | NA | + | 0.08 | + | NA | NA | NA | NA |
| haem_all_sites_model1 | 8 | -1089.1 | 2194.2 | 0.00 | 0.26 | 1.37 | + | + | NA | 0.10 | NA | + | NA | NA | + |
| haem_all_sites_model2 | 7 | -1090.4 | 2194.8 | 0.60 | 0.19 | 1.37 | + | NA | NA | 0.10 | NA | + | NA | NA | + |
| haem_all_sites_model3 | 9 | -1088.8 | 2195.8 | 1.52 | 0.12 | 1.33 | + | + | + | 0.10 | NA | + | NA | NA | + |
| plas_all_sites_model1 | 7 | -1421.4 | 2856.8 | 0.00 | 0.14 | 0.68 | + | NA | + | 0.06 | NA | NA | NA | NA | + |
| plas_all_sites_model2 | 6 | -1422.4 | 2856.9 | 0.09 | 0.13 | 0.74 | + | NA | NA | 0.06 | NA | NA | NA | NA | + |
| plas_all_sites_model3 | 8 | -1420.6 | 2857.2 | 0.41 | 0.11 | 0.67 | + | NA | + | 0.05 | NA | NA | + | NA | + |
| plas_all_sites_model4 | 8 | -1421.2 | 2858.4 | 1.60 | 0.06 | 0.68 | + | + | + | 0.06 | NA | NA | NA | NA | + |
| plas_all_sites_model5 | 8 | -1421.2 | 2858.4 | 1.64 | 0.06 | 0.67 | + | NA | + | 0.06 | NA | + | NA | NA | + |

| | | | | | | | | | | | | | | | |
|---|---|---|---|---|---|---|---|---|---|---|---|---|---|---|---|
| plas_all_sites_model6 | 7 | -1422.2 | 2858.5 | 1.72 | 0.06 | 0.74 | + | + | NA | 0.06 | NA | NA | NA | NA | + |
| plas_all_sites_model7 | 7 | -1422.3 | 2858.6 | 1.76 | 0.06 | 0.74 | + | NA | NA | 0.06 | NA | + | NA | NA | + |
| plas_all_sites_model8 | 8 | -1421.4 | 2858.8 | 1.98 | 0.05 | 0.67 | + | NA | + | 0.06 | + | NA | NA | NA | + |
| leuco_all_sites_model1 | 5 | -1204.2 | 2418.4 | 0.00 | 0.08 | 1.32 | + | NA | + | 0.03 | NA | NA | NA | NA | NA |
| leuco_all_sites_model2 | 6 | -1203.3 | 2418.7 | 0.26 | 0.07 | 1.32 | + | NA | + | 0.04 | NA | + | NA | NA | NA |
| leuco_all_sites_model3 | 4 | -1205.5 | 2419.0 | 0.59 | 0.06 | 1.38 | NA | NA | + | 0.03 | NA | NA | NA | NA | NA |
| leuco_all_sites_model4 | 6 | -1203.7 | 2419.4 | 0.92 | 0.05 | 1.28 | + | NA | + | 0.03 | + | NA | NA | NA | NA |
| leuco_all_sites_model5 | 5 | -1204.7 | 2419.5 | 1.04 | 0.05 | 1.36 | NA | + | + | 0.03 | NA | NA | NA | NA | NA |
| leuco_all_sites_model6 | 7 | -1202.8 | 2419.6 | 1.19 | 0.04 | 1.28 | + | NA | + | 0.04 | + | + | NA | NA | NA |
| leuco_all_sites_model7 | 6 | -1204.0 | 2420.0 | 1.59 | 0.04 | 1.32 | + | + | + | 0.03 | NA | NA | NA | NA | NA |
| leuco_all_sites_model8 | 7 | -1203.1 | 2420.2 | 1.79 | 0.03 | 1.33 | + | + | + | 0.04 | NA | + | NA | NA | NA |
| triple_all_sites_model1 | 7 | -1548.6 | 3111.2 | 0.00 | 0.16 | -0.40 | + | NA | + | 0.08 | NA | NA | NA | NA | + |
| triple_all_sites_model2 | 6 | -1550.0 | 3112.0 | 0.81 | 0.10 | -0.33 | + | NA | NA | 0.08 | NA | NA | NA | NA | + |
| triple_all_sites_model3 | 8 | -1548.2 | 3112.4 | 1.18 | 0.09 | -0.40 | + | + | + | 0.08 | NA | NA | NA | NA | + |
| triple_all_sites_model4 | 8 | -1548.3 | 3112.7 | 1.43 | 0.08 | -0.40 | + | NA | + | 0.08 | NA | NA | + | NA | + |
| triple_all_sites_model5 | 8 | -1548.4 | 3112.8 | 1.55 | 0.07 | -0.40 | + | NA | + | 0.09 | NA | + | NA | NA | + |
| triple_all_sites_model6 | 8 | -1548.5 | 3113.0 | 1.80 | 0.06 | -0.42 | + | NA | + | 0.08 | + | NA | NA | NA | + |
| triple_all_sites_model7 | 7 | -1549.6 | 3113.2 | 1.92 | 0.06 | -0.33 | + | + | NA | 0.08 | NA | NA | NA | NA | + |

**Table 3** – Summary of lineages by sampling site. The lineage beginning with "H" is *Haemoproteus*, lineages starting with "P" are *Plasmodium*, and lineages that start with "L" are *Leucocytozoon*.

| Lineage | Åsen | Öved | Revinge | Total |
|---|---|---|---|---|
| H_PARUS1 | 12 | 59 | 587 | 630 |
| P_BT7 | 0 | 3 | 36 | 39 |
| P_SGS1 | 0 | 1 | 1 | 2 |
| P_TURDUS1 | 1 | 13 | 49 | 63 |
| P_Unknown | 1 | 1 | 8 | 10 |
| L_BT2 | 0 | 2 | 2 | 4 |
| L_PARUS12 | 0 | 4 | 9 | 13 |
| L_PARUS14 | 0 | 2 | 12 | 14 |
| L_PARUS18 | 0 | 3 | 8 | 11 |
| L_PARUS21 | 0 | 2 | 2 | 4 |
| L_PARUS22 | 0 | 0 | 1 | 1 |
| L_PARUS4 | 0 | 9 | 18 | 27 |
| L_PARUS74 | 0 | 0 | 4 | 4 |
| L_COINFECTION | 0 | 5 | 27 | 32 |

**Table 4 –** Predictors for *Haemoproteus*, *Plasmodium*, and *Leucocytozoon* infection from top performing models ($\Delta$AIC ≤ 2) fit to the Revinge-only dataset. Model-averaged logistic regression coefficients on the log-odds scale (Log-odds), associated standard errors (SE), and p-values (p) are shown. Summed Akaike weight across the top-performing models in which each predictor was included.

| Parasite | Predictor | Log-odds | SE | p | Weight |
|---|---|---|---|---|---|
| *Haemoproteus* | (Intercept) | 1.39 | 0.11 | **< 2.2E-16** | NA |
| *Haemoproteus* | Age = Older | 1.12 | 0.19 | **< 2.2E-16** | 1 |
| *Haemoproteus* | Sex = Male | 0.11 | 0.14 | 4.33E-01 | 0.54 |
| *Haemoproteus* | Year | 0.11 | 0.01 | **< 2.2E-16** | 1 |
| *Haemoproteus* | Age = Older:Year | 0.04 | 0.02 | **4.50E-02** | 1 |
| *Haemoproteus* | Recaptured = TRUE | 0.12 | 0.24 | 6.14E-01 | 0.39 |
| *Plasmodium* | (Intercept) | 0.61 | 0.09 | **< 2.2E-16** | NA |
| *Plasmodium* | Age = Older | 0.79 | 0.13 | **< 2.2E-16** | 1 |
| *Plasmodium* | Sex = Male | 0.30 | 0.11 | **9.52E-03** | 1 |
| *Plasmodium* | Year | 0.05 | 0.01 | **2.80E-06** | 1 |
| *Plasmodium* | Recaptured = TRUE | -0.02 | 0.09 | 8.37E-01 | 0.15 |
| *Plasmodium* | Age = Older:Year | -0.00 | 0.01 | 8.43E-01 | 0.15 |
| *Leucocytozoon* | (Intercept) | 1.32 | 0.11 | **< 2.2E-16** | NA |
| *Leucocytozoon* | Sex = Male | 0.29 | 0.14 | **3.22E-02** | 1 |
| *Leucocytozoon* | Year | 0.04 | 0.01 | **4.14E-05** | 1 |
| *Leucocytozoon* | Age = Older | 0.12 | 0.16 | 4.42E-01 | 0.60 |
| *Leucocytozoon* | Age = Older:Year | -0.01 | 0.01 | 6.52E-01 | 0.27 |
| *Leucocytozoon* | Recaptured = TRUE | 0.015 | 0.08 | 8.51E-01 | 0.10 |
| Triple infection | (Intercept) | -0.44 | 0.08 | **2.00E-07** | NA |
| Triple infection | Age = Older | 0.76 | 0.11 | **< 2.2E-16** | 1 |
| Triple infection | Sex = Male | 0.25 | 0.10 | **1.20E-02** | 1 |
| Triple infection | Year | 0.09 | 0.01 | **< 2.2E-16** | 1 |
| Triple infection | Age = Older:Year | -0.01 | 0.01 | 5.75E-01 | 0.40 |
| Triple infection | Recaptured = TRUE | 0.04 | 0.12 | 7.04E-01 | 0.26 |
